# Size switchable DNA origami structure enabled by dynamic crossovers

**DOI:** 10.1101/2025.02.07.637188

**Authors:** Xueqiao Li, Rui Gao, Xiaoxue Wang, Wenna Wu, Huajie Liu, Tao Zhang

**Affiliations:** Department of Applied Chemistry, School of Chemistry and Chemical Engineering, Yantai University, Yantai 264005, China; School of Chemical Science and Engineering, Shanghai Research Institute for Intelligent Autonomous Systems, Key Laboratory of Advanced Civil Engineering Materials of Ministry of Education, Tongji University, Shanghai, 200092, China

**Keywords:** DNA origami, crossover, i-mo9f, DNA hairpin, size-switchablity

## Abstract

DNA nanostructures are formed by many DNA double helices connected via crossovers. If multiple DNA double helices are connected by crossovers at different base positions (different rotational angles), these DNA double helices are stacked together to obtain desired two- or three-dimensional DNA structures. Here we inserted i-motif or hairpin motifs in crossover regions of one layer DNA origami structure and employed dynamic DNA element to implement size-switchablity between expansion and contraction. By changing pH or adding complementary strands to regulate interspacings between DNA double helixes, multiple crossover replacements realize the dynamic size changes between 68 nm and 150 nm as characterized by atomic force microscopy. We further demonstrated dynamic sizes changes enabled or inhibited FRET (fluorescence resonance energy transfer) signals between Cy3 and Cy5 and the controlled distances between protein molecules. The results reveal the feasibility of fabricating size-switchable DNA origami nanostructures via crossover redesign, demonstrating their potential in nano-engineering, proximity enabled chemical reacitons, as well as for biomedical applications.

## Introduction

In the native four-stranded DNA intermediate Holliday junction,^1,2^ two DNA duplex were connected via one crossover linkage. Resembling Holliday junction, DNA nanostructures are usually built by crossover linked multiple DNA helix bundles.^3^ Hence the rigid double stranded DNA and crossover connections can be regarded as two primary structural elements in structural DNA nanotechnology. In particular, owing to the helical feature of double-standed DNA, crossover positions determine the attachement angle of each newly-added helix. Taking DNA origami technique as an example, the double crossover connection position can align DNA helix in parallel to form one dimensional single-layered DNA origami^4^ or pack DNA helix in cubic or honeycomb lattices^5^ to fold three dimensional solid DNA structures. In other words, crossover positions, DNA helix numbers, and DNA helix lengths can determine the mophorlogies of DNA nanostructures. Over the past decades, DNA has proven to be a powerful tool for assembling customized nanostructures, with several assembly strategies have been developed.^6–8^ Mostly have focused on designing the hybridization rules between different segments among DNA oligonucleotides. Firstly, the well-defined duplex DNA follows strick nitrogenous base pairing rule of adenine (A) = thymine (T) and cytosine (C) expandable ≡ guanine (G). As a low energy favored exoergic reaction, the hybridization can be conducted by properly controlled temperature ramps. Secondly, the mature nucleic acid chemistry permitted orthogonal sequence design^9^ and solid-state based DNA synthesis with relatively low price, which lay the material basis for feasible lab experiments.

Moreover, if a special crossover, e.g. embedded with DNA spacer of various lengths, is introduced as a pivot joint to link DNA substructure, switchable DNA actuators,^10^ robotic DNA arms,^11^ as well as complex DNA structures^12^ can be obtained. For instance, while the linkage in shaple-complementary nanoswitch contains zero base spacer^13^, in the reconfigurable 3D plasmonic metamolecules, two substructures are connected via 8-nt region of scaffold to render a flexible linkage^14^. In the electric field controlled robotic arm, the “arm” is connected to the plate via an asymmetric crossover of 3 and x-nt (x=0, 3, 4, 7, 13, or 23) scaffold spacer,^15^ allowing for rotational movement according to electric field as well as the winding up as a mecocular torsion spring. Many biomedical or photonic applications have employed such switchable DNA structural motif to realize a dynamic control.^16,17^ As crossover is key element to link neighboring duplexes, we envision that by replacing multiple crossovers in DNA origami with a dynamically extensile and contractile unit, the distances between each DNA double helix can be regulated. Pecularly, in two-dimensional one-layer DNA origami sheet, the crossovers in every neighbouring duplx are replaced and the transformation can be accumulated to manyfold.

There are well-characterized DNA-based intramolecular extensile and contractile units that are usually encoded with intrastrand self-complementary sequences such as stem-loop hairpin and i-motif.^18^ While the formation of intramolecular structures shortened the end-to-end distance, it can be reversibly opened and extended to full distance by hybridizing to its complementary strands. In addition, the C-rich DNA molecules i-motif is pH-responsive which can form quadruplex stabilized by intercalated, hemi-protonated cytosine-cytosine base pairs in a narrow pH range of 4.0-4.5.^19^ To realize such robust morphology transformation that responsive to pH or target strand, it requires only appropriate sequence design with certain strand purity. Previous studies have not only examined different operative conditions for opening and closing of the stem-loop hairpin or i-motif, but have also positioned these extensile and contractile units on nanostructures to implement actuational movements. Such as hairpin DNA was used as a single stranded tether to hold tweezer arms within a certain radius and minimize side reactions so to improve the tweezing efficiency.^20^ Liu et al. incorporated i-motif unit in a DNA tetrahedron framework and employed its pH-sensitivity to develop a size-adaptive tetrahedron according to pH variations in single synaptic vesicles.^21^ M. Majikes et al. designed parallel i-motif leash to seam up two half-origami.^22^ The formation of i-motif self-intercalating secondary structure shortened the end-to-end distance hence realized a large scale motion between two half-origami. We here replaced all crossovers in one-layer DNA origami with extensile and contractile units and demonstrated a size-switchable DNA structure.

## Results and Discussion

We chose one-layer DNA origami structure 32 helix bundle (32HB) as a prototypical structure for crossover redesign. To implement customerized sequence design, 5 or 6 crossovers to connect every neighboring duplexes are via staples while the scaffold DNA is routed in a zig-zag style with 42-nt single stranded endcap loop. We replaced all crossovers in the upper 16HB (green) with dynamic units and keep original crossover design in the lower 16HB (gray) as static control. (Figure 1, Figure S1). Crossover staples in the expandable region were inserted self-complementary sequences such as i-motif or DNA hairpin (Figure 1a). We used 25-nt i-motif sequence -AA CCC TAA CCC TAA CCC TAA CCC AA- and DNA hairpin sequence -T GCG ACC GT TTTTTTT ACG GTC GC T-. In contracted states, these dynamic units functioned as protruded joints to link parallel duplexes. When hybridizing to their respective complementary strands, secondary structures are opened and extended to a double strands thus expands the duplex-duplex distances. Comparing to the characterized inter-duplex spacing in DNA origami of 2.7 nm,^23^ the fully extended duplex spacing with dynamic crossovers should be 8.5 nm (25 base pairs). With 16 neighboring duplexes linked by dynamic crossovers, the fully expansion should be able to enlarge the structural length from 40.5 nm to 157.5 nm.

**Figure 1.**
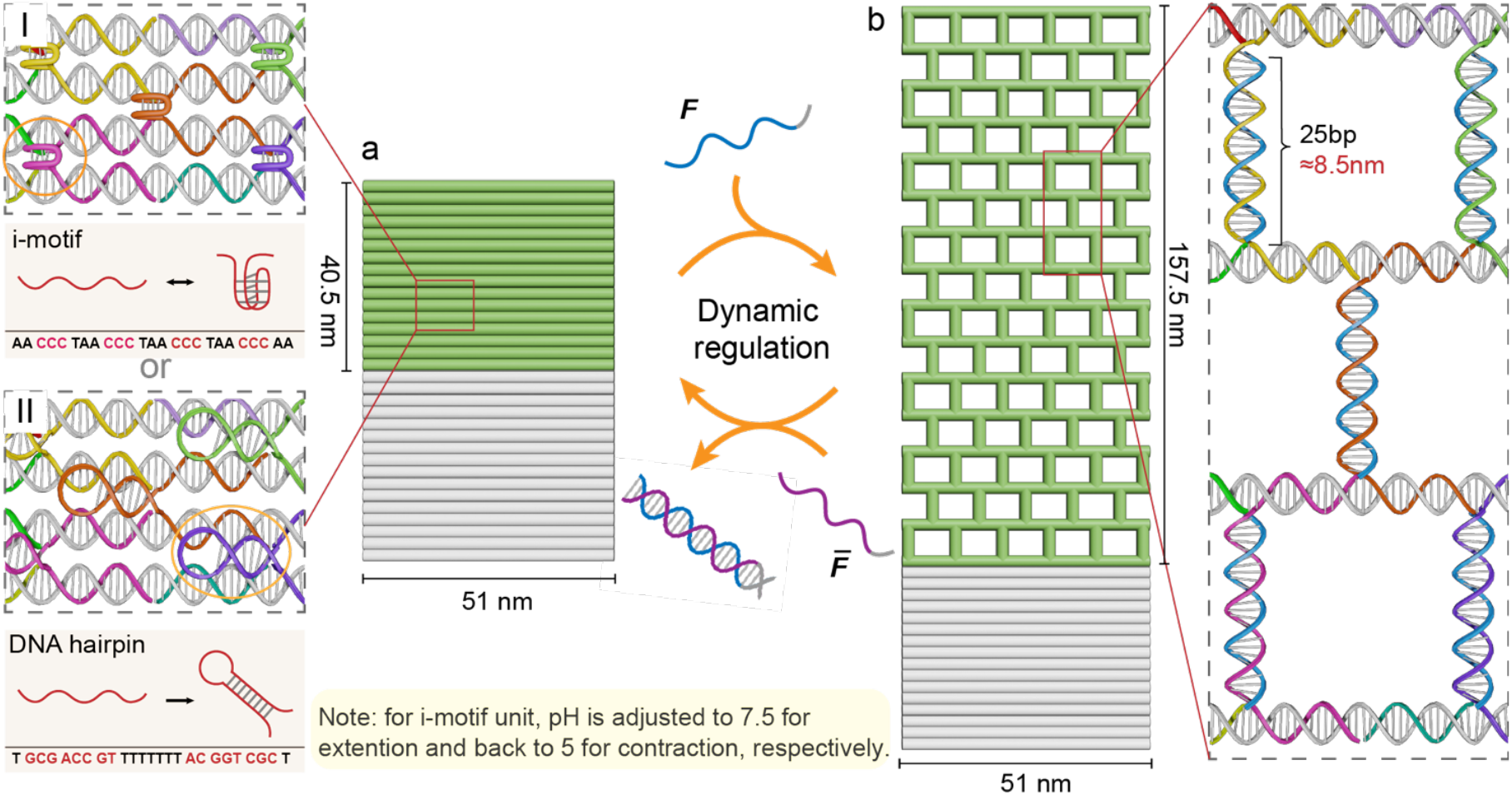
Schematic illustration of size-switchable DNA origami nanostructures. (a) One layer DNA origami in the contracted state. It consists of two parts, the upper part (green) is the crossover replaced expandable structure, and the lower part (gray) is DNA structure for control. There are two kinds of responsive crossover units inside the expandable part: i-motif or DNA hairpin. (b) The extended state of DNA nanostructure when complementary strands F opened the secondary structure to form a duplex which theoretically enlarge the structure to about two times bigger. The removal of F by fuel strand FJ reverse the structure to contracted state (F/FJ pairs are only symbol but the sequence for i-motif and hairpin are different). For i-motif enabled extension, the pH is accordingly adjusted from 5 to 7.5. When contracting the structure, fuel strand FJ is added to remove F strand and pH is titrated to 5 again. The designed structure width is about 51 nm with the expandable part have the size change capacity from 40.5 nm to 157.5 nm.

The origami folding were carried out with standard procedure using mini-M13 scaffold DNA and 10x times staple strands in 1x TAE (Tris Acetate-Ethylenediaminetetraacetic acid buffer), 10mM MgCl_2_ annealed under temperature ramp 65°C-25°C for 2 hrs (Table S1-3). To open the hairpin or i-motif, the complementary strands *F* were added and thermally annealed from 55°C to 25°C for 2hrs (Figure S2-3). The adding of fuel strands remove the complementary strands and drive the structure to contracted states. For DNA origami nanostructure comprising i-motif, the pH was accordingly adjusted from 5 to 7.5 during extension to double stranded DNA and backwards to 5 for i-motif formation.

Next, we verified DNA nanostructures folding and its size-switchablibility using agarose gel electrophoresis and atomic force microscopy (AFM) (Figure 2). Figure 2a shows the agarose gel electrophoresis (1%, 65 V, 1x Tris Acetate-Ethylenediaminetetraacetic acid buffer, 2hrs) images of 32HB containing i-motif replaced crossovers, and the static origami structure 16HB as control. The gel bands showed that the contracted 32HB have similar mobility with the control structure at pH=5. When all i-motif quadruplex were opened, the extended structures showed delayed mobility than the control, indicating a larger structure at pH=7.5. AFM characterization verified the sizes differences, 32HB at pH=5 showed the static control section 16HB has a length of about 50.1 nm and the rest i-motif replaced crossover section is about 68.7 nm (Figure 2b). We think i-motif junctures occupied considerable spaces and gave rise to larger inter-duplex spacings. Ager opening the i-motif with its complementary strands, the extended structure at pH=7.5 showed a length of about 151.3 nm, more than two-times larger (Figure 2c). For structures of hairpin replaced crossovers, the hairpins were opened by adding its complementary strands to reach extended state. Additionally adding fuel strands F can remove strand F and reverse the structure to contracted state. Agarose gel electrophoresis showed contracted and extended 16HB caused different mobilities for 32HB. Though the mobility differences are subtle, it implies the size differences. AFM images gave further evidences about the size-switchability. The measured length of static 16HB is about 50.8 nm and the 16HB containing hairpin replaced crossovers in contracted state is about 67.7 nm, 17 nm larger than the static control (Figure 2e). Again, we think the joints using hairpin structure causes the size differences comparing to structures made of conventional crossovers and hence both kinds of structures in the contracted state are larger than calculated value. For extended structures of both replaced crossovers, it presents mesh-shaped structures due to the fully opened hairpin or quadruplex, verifying the crossover redesign could extend the one-layer origami structure to more than 2 times larger.

**Figure 2.**
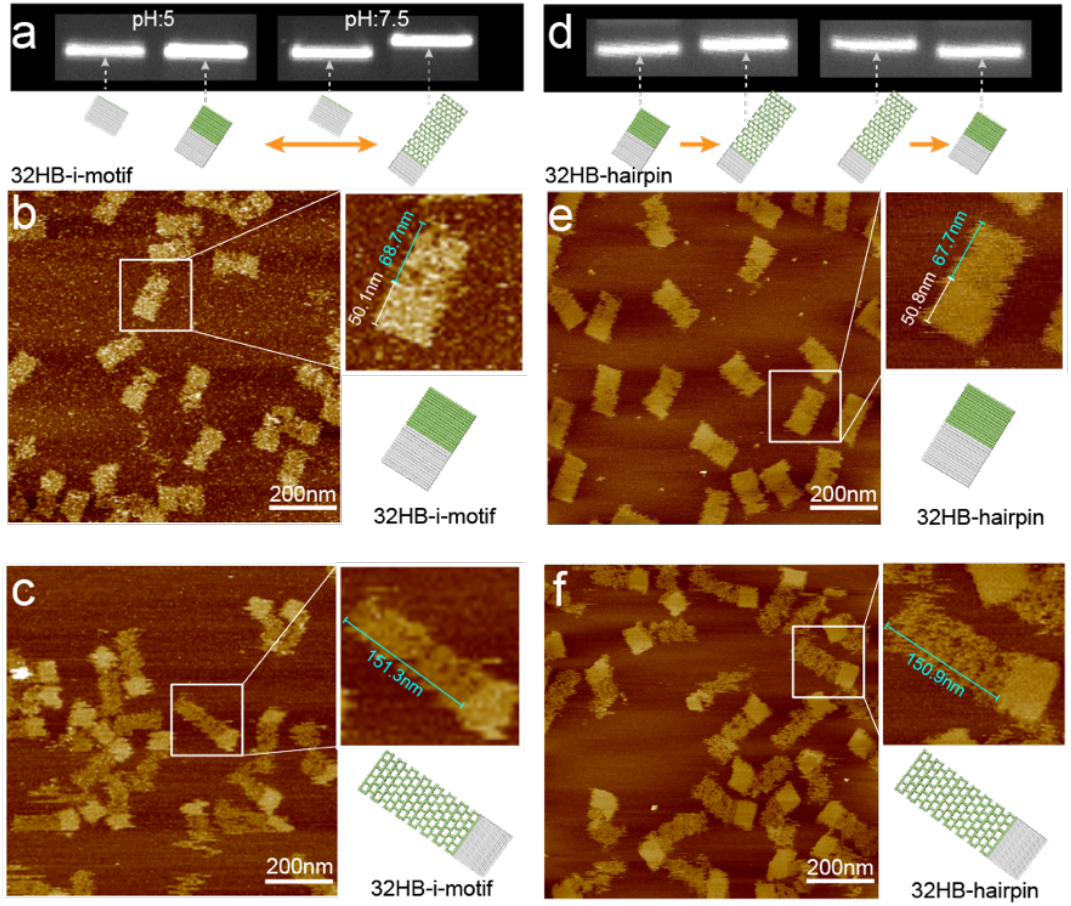
Characterization of the size switchable DNA origami nanostructures. (a) Agarose gel electrophoresis (1%, 65 V, 1x Tris Acetate-Ethylenediaminetetraacetic acid buffer buffer, 2hrs) to characterize i-motif enabled extended and contracted structure. Comparing to the control structure, the extended structures have much delayed mobility. (b) Atomic force microscopy (AFM) characterization showed the structure of normal crossovers is 50.1 nm while the contracted i-motif section is 68.7 nm. (c) After extend the C-quadruplex to duplex, structure is extended to 151.3 nm. (d) Agarose gel electrophoresis to characterize hairpin enabled expansion and contra5on. The expanded structure showed delayed mobility comparing to the unextended 32HB origami structure. (e) AFM image showed showed the structure of normal crossovers is 50.8 nm while the contracted hairpin section is 67.7 nm. (f) The fully extended structure reached 150.9 nm after opening hairpin structure.

To test the size-switchability, we labeled FRET (fluorescence resonance energy transfer) dye pairs on neighbouring duplex and investigated the reversible structural changes. It is known that photon energies can be transferred from one excited fluorophore (donor) to another (acceptor) when the donor and acceptor of fluorescent molecules are in close proximity to each other. When the structure is in contracted state, it enabled the energy transfer from Cy3 to Cy5 and the emission intensities of donor dye decreased while the emission intensities of acceptor dyes increased. When the structure is extended and the spacing of two dyes is beyond FRET distances, the energy transfer is inhibited (Figure 3a). Excited with 547 nm, we collected the emission of Cy5 and explored the influences of expansion and contraction of 32HB to the luminescence intensities of acceptor dyes. Figure 3c shows the fluorescence spectras of Cy5 on 32HB containing i-motif replaced crossovers for multi changing rounds. When it was in the contracted state at pH 5.0 (32HB-1,3 in Figure 3c), the energy transfer efficiency is high as indicated by the emission intensities of Cy5. When in the extended state at pH 7.5 (F-32HB-2,4 in Figure 3c), FRET was blocked as shown in the decreased Cy5 fluorescence intensities. The i-motif enabled reversible size changes enabled the alternative FRET signals changes of Cy3 to Cy5. However, we also see the signal damping due to the side reactions and accumulated fuels. The 32HB containing hairpin replaced crossovers showed similar tendency. In the folded hairpin structure, the one-layer DNA origami is in a contacted state which enabled the FRET singnal as shown in Figure xx. The opened hairpin structure to duplex extended the structure and blocked the energy transfer from Cy3 to Cy5. By comparing maximum emission at 665 nm of the two nanostructures (Figure 3e), structures containing i-motif replaced crossovers showed more pronounced signals changes but also decreased faster comparing to hairpin crossovers. The reversible opening and folding of i-motif quadruplex, it requires buffer exchanges which contains process of multi-rounds centrifugation and cause damages and samples loss. Nevertheless, the FRET measurements supported our crossover redesign enabled size-switchability.

**Figure 3.**
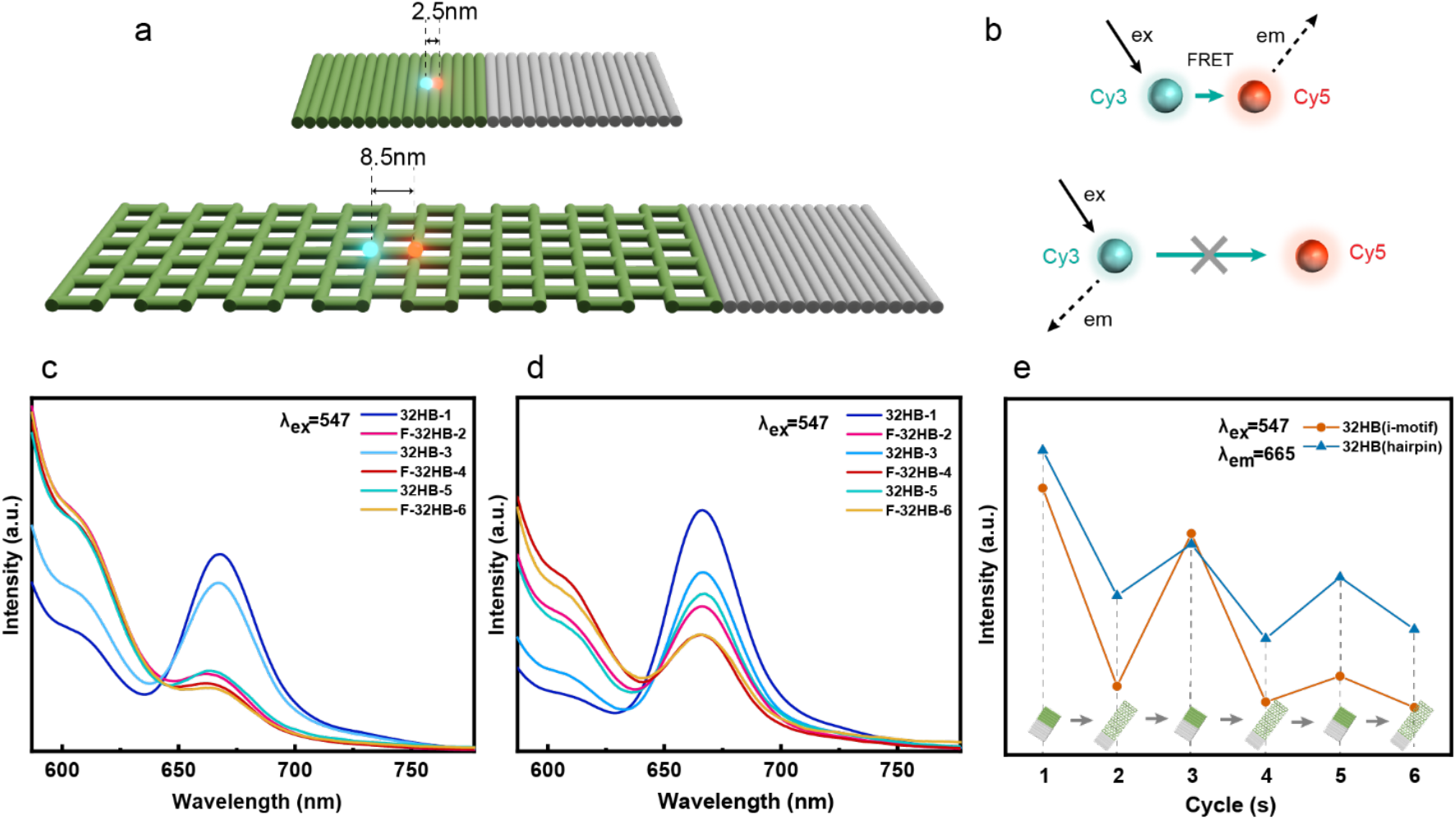
The fluorescence resonance energy transfer (FRET) on expandable DNA origami nanostructures. Schematic diagram of expandable one-layer DNA origami labelled with fluorescent molecules. As the structure extended, it lead to larger distance between the fluorescent molecules (e.g. Cy3 & Cy5). (b) Schematic representation of FRET effect. When two fluorescence molecules are close to each other, photon energies can be transferred from one excited fluorophore (Cy3) to the other (Cy5). Larger spacings will inhibit the FRET effect. (c) Fluorescence emission profiles of DNA structure with responsive unit being i-motif. 1-6 are the number of structural transitions, 32HB is the structurally contracted state, and F-32HB is the structurally extended state. (d) Fluorescence emission profiles of nanostructural transitions with DNA hairpin as the responsive unit. (e) Comparison of fluorescence intensi5es at the wavelength 665 nm for six 5mes of nanostructure transistions.

In addition, interspacings are crucial to regulate multienzymatic cascade reactions and the contraction and expansion allows to construct functional arrays with dynamic spacings. As a proof of concept, we assembled four sets of biotin modified DNA strands located at selected positions on 32HB made of hairpin crossvers and achieved avidin-avidin distances changing along with structural scalability (Figure 4a). AFM images verified four conjugated streptavidin on biotin sites with interspacing about 15.8 nm (Figure 4b, c), and enlarged distances to about 27.5 nm between them resulted from structural expansion. This demonstrated the capacity to facilitate programmal dynamic assemblies on the nanoscale and to study the complex molecular interactions.

**Figure 4.**
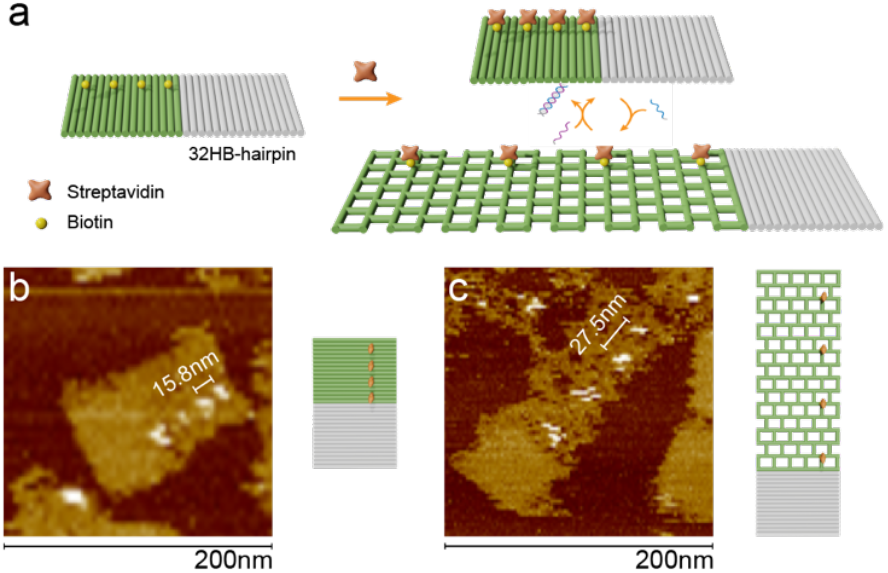
Assembly of proteins with dynamic spacings on expandable DNA origami nanostructures. (a) Schematic representation of the surface-modified biotin (yellow)-streptavidin (orange) on the 32HB. (b) Atomic force microscopy (AFM) characterization of four streptavidin conjugated to biotin molecules on 32HB in the contracted state. (c) AFM characterization of four streptavidin conjugated to biotin molecules on 32HB in the extended state.

## Conclusion

In summary, we explored a means to design expandable and contractable DNA origami nanostructures by inserting dynamic units, i-motif or DNA hairpin, in crossovers that connect neighboring duplexes. AFM characterization showed the expansion can enlarge from 68 nm to 150 nm. We further showed the size switchability with FRET signals and dynamic array of protein assemblies. In general, size changes not only can switch spacings of dynamic arrays but also are crucial to reach complex shape transformations. For instance, in the antitumor drug delivery area, there is paradox of between circulation time and particle sizes.^24–26^ While smaller drug carriers have a better penetration, it also showed faster clearances due to the high fluid pressure inside the tumor. Larger carriers are capable of a longer circulation and tumor retention but it is hard to penetrate into the dense tumor matrix. DNA structures have been demonstrated in many places for accurate drug carrier design. Here the crossover redesign will allow to apply reversible size-changing methodology for intelligent size control so to reach the critical point of equilibrated penetration and retention. Hence a contracted small carrier with targeting molecules and drug payload can be delivered to tumor region and expanded its size to minimize eliminatio rates. Thus, we envision this dynamically swelling and shrinking nanostructures will have great potential for applications in optical and biochemical areas.

## Acknowledgements

This work is financially supported by the Science and Technology Committee of Shanghai Municipality (2022-4-ZD-03), the Shanghai Pilot Program for Basic Research, the Fundamental Research Funds for the Central Universities, and the Natural Science Foundation of Shandong Province (ZR2020QB163). T.Z. thanks the generous funding support from the Taishan scholars’ program of Shandong province (tsqn201909083), China.

## Declaration of Interests

The authors declare no conflict of interest.

## Notes

### Competing Interest Statement

The authors have declared no competing interest.

