## Supplementary material for "Size switchable DNA origami structure enabled by dynamic crossovers": SI

### Supporting Information

**Content**


**1 Materials and methods**

**1.1 Materials**

**1.2 Synthesis of stretchable DNA origami nanostructures**

**1.3 Structural scaling transformations**

**1.4 Agarose gel electrophoresis assay**

**1.5 AFM characterization**

**1.6 Fluorescence spectral analysis**

**1.7 Protein localization assembly**

**2 Supporting tables**

**2.1 tables S1-S4**

**3 Supporting figures**

**3.1 figures S1-S5**

**4 Additional references**

#### 1. Materials and methods

##### 1.1 Materials

All oligonucleotides were synthesized and purified by Tsingke Biotechnology Co., Ltd. Oligonucleotide were purified by HPLC. Chemicals were directly used without further purification. All solutions were prepared by ultrapure water (resistance = 18.2 MΩ*cm), which was obtained through a Millipore Milli-Q ultrapure water system (Milli-Q Reference, FR). Electrophoresis apparatus (SH-DYC10V, SH-SP10H) and gel imager (SH-510) were purchased from Hangzhou Shenhua Tech. Co. Ltd. PCR gene amplifier was purchased from Suzhou Dongsheng Xingye Scientific Instrument Co. Ltd. The microcentrifuge was purchased from Eppendorf, Germany.

##### 1.2 Synthesis of stretchable DNA origami nanostructures

The two DNA origami structures mentioned in this study consisted of 32 helical bundles (Fig. S1). The structure design was performed using caDNAno software and corrected according to the analysis results obtained from the CanDo platform. The two structures consisted of DNA origami framework scaffold chains (M13mp18s), staple strands in the normal origami section (Table S1), staple strands in the retractable section (32HB-i-motif: Table S2, 32HB-hairpin: Table S3), and dynamically converted complementary strands (Table S4) were composed. The DNA strands were mixed in buffer at a molar ratio of 1:10 (scaffold: staple), the buffer used for 32HB-i-motif was 1× sodium acetate-12 mM $Mg^{2+}$ (50 mM sodium acetate, 1 mM EDTA, 12 mM $MgCl_2$) at pH=7.5 and pH=5, and the buffer used for 32HB-hairpin was 1× TAE-12 mM $Mg^{2+}$ (40 mM Tris base, 20 mM Glacial acetic acid, 1 mM EDTA, 12 mM $MgCl_2$). The origami structural monomer concentration during assembly was 10 nM, and annealing was performed from 65°C to 20°C using a PCR device for 2 h (65°C for 15 mins; 65 to 20°C, -0.2°C/25s; 8°C). Subsequent enrichment and purification of the structure is achieved using ultrafiltration purification[1] and PEG purification[2], with the possibility of buffer replacement in the process. Purification by ultrafiltration: the sample was placed in a 100 kp ultrafiltration tube and buffer (1× sodium acetate-10 mM $Mg^{2+}$ buffer/1× TAE-10 mM $Mg^{2+}$ buffer, depending on the structure) was added to 400 μL. The sample was centrifuged at 5000 rpm for 8 min, and then the buffer was added to repeat the centrifugation for three times. After that, inverted ultrafiltration tube was centrifuged for 2 min to obtain the purified sample. PEG purification: the sample and PEG buffer (15% (w/v) PEG-8000, 2× TAE, 500mM NaCl, 20mM $MgCl_2$) were mixed well at 1:1, (13000rpm, 32min), the supernatant was removed, and the bottom precipitate was added to the buffer (determined according to the structure) by shaking for 12h (700rpm, 30°C). The monomer concentration obtained after purification was 20nM~50nM, and stored at -20°C for the next step.

##### 1.3 Structural scaling transformations

Each stretching change of this stretchable nanostructure was accomplished in PCR (thermal annealing:

55°C-25°C, 2h). For the stretching changes we used the ultrafiltration (as above) method to remove excess DNA strands and perform buffer replacement. To realize the 32HB-i-motif structure from contracted to stretched state, the buffer of the contracted sample was firstly replaced with 1× sodium acetate-10mM $Mg^{2+}$ buffer at pH=7.5, and after that, the structure in the stretched state was obtained by adding 10-fold complementary strand i-motif-a and thermally annealing it for 2h. The 32HB-i-motif structure was then changed to the contracted state by replacing the buffer with 1× sodium acetate-10mM $Mg^{2+}$ buffer at pH=5 and adding 10-fold complementary chain i-motif-b for 2h of thermal annealing. To realize the change of 32HB-hairpin structure from contracted to stretched state, it can be obtained by directly adding 10-fold complementary strand hairpin-a to the sample by thermal annealing for 2h. Afterwards, the excess DNA strands are removed using ultrafiltration, and the 32HB-hairpin structure can be changed from a stretched to a contracted state by adding 10-fold complementary strand hairpin-b heat annealed for 2h.

**1.4 Agarose gel electrophoresis assay**

The assembled nanostructures were characterized using agarose gel electrophoresis (Agarose Gel Electrophoresis, AGE). A 1% agarose gel was used in this experiment and stained with 1% Ethidium Bromide (EB) with electrophoresis conditions of 65v, 2h. Samples of 32HB-i-motif changing from contraction to elongation and from elongation to contraction again were subjected to gel electrophoresis with 1% agarose gel in 1× sodium acetate-10 mM $Mg^{2+}$ buffer at pH=7.5 and pH=5, respectively, along with the plain origami structure 16HB (Fig. 2a). Samples of 32HB-hairpin that changed from contraction to elongation and from elongation to contraction again were subjected to gel electrophoresis with a 1% agarose gel in 1× TAE-10 mM $Mg^{2+}$ buffer (Fig. 2b). After electrophoresis, the agarose gel was transferred to a UV imager for observation (UV transmission). Comparison of the band positions verified the telescopic reversibility of the two DNA origami structures.

**1.5AFM characterization**

AFM imaging was carried out on a Dimension FastScan AFM (Bruker Corp.) with ScanAsyst Fluid + probes in fluid at the ScanAsyst mode. The sample solution (1 nM ,5 μL) was deposited onto a freshly cleaved round mica disk and allowed to stay for 3 min. After removing the remaining solution, the mica sheets were washed with buffer (1× sodium acetate-10mM $Mg^{2+}$ /1× TAE-10mM $Mg^{2+}$). Finally, 15 μL of Buffer were deposited onto the mica disk and 15 μL of Buffer onto the AFM tip before imaging。

**1.6 Fluorescence spectral analysis**

Fluorescence detection verified the FRET effect of the structures in the contracted/stretched state. The assembly of DNA origami nanostructures containing fluorescent molecules was first carried out (the steps were the same as for the assembly of DNA origami nanostructures and were performed under light protection). After purification by ultrafiltration, the concentration of the origami nanostructures was determined by ultra-micro UV-visible spectrophotometer after dilution with appropriate buffers (1× sodium acetate-10 mM $Mg^{2+}$ buffer/1× TAE-10 mM $Mg^{2+}$ buffer, depending on the structure).Afterwards, 1 ml (concentration of about 5 nM) of the sample was placed in a quartz cuvette and the fluorescence emission profile was obtained by fluorescence spectrometer (F-4700) detection with an excitation light wavelength of 547 nm. The 32HB-i-motif and 32HB-hairpin were subjected to six structural telescopes, each purified using ultracentrifugation with simultaneous buffer replacement. The concentration of origami nanostructures was measured using an ultra-micro UV-visible spectrophotometer, followed by fluorescence detection again. The curve was fitted according to the concentration of origami nanostructures to obtain a fluorescence curve for the six times of structure stretching.

##### 1.7 Protein localization assembly

First, assembly experiments of 32HB-hairpin with aligned biotin were performed (the steps were the same as those for the assembly of DNA origami nanostructures). After purification by ultrafiltration, it was diluted to 5 nM with 1×TAE-10 mM $Mg^{2+}$ buffer. after that, streptavidin (solvent: 1×TAE-10 mM $Mg^{2+}$ buffer) was added to the samples at a ratio of 10:1 (streptavidin: biotin), and the reaction was carried out for 12h at 25°C.

#### 2 Supporting tables

##### 2.1 tables S1-S4

Tables S1

| Name | Sequence (5' - 3') |
|---|---|
| core1 | TTAGCAACAGTAGGGCTTAAGCCAGAGGGGGTA |
| core2 | GAATTACAAATTCTTACCAGGATAGCGTCC |
| core3 | CCTTTAGAAAAAGCCTGTTTAATATTCATTGA |
| core4 | TAACTAAATAAGAATAAACAACAGTTCAGAAA |
| core5 | ATGCAAATCCAATCGCTGAAATACCGACCGTGAAAATCAGGTCT |
| core6 | ATACGTTTACCAGACGACGTCCAGACGACGACAATAGA |
| core7 | GACTGTATAAAGCCAACGCTATTAAGACGCTGAGAA |
| core8 | AATAGGCATAGTAAGAGCATAATGCAGAACGCGCGAC |
| core9 | CATAAGTATCATATGCGTTATTATCAAAATCATAGG |
| core10 | ATCCATGCAGATACATAACGGATAAGTCCTGAACAACTA |
| core11 | TTAACCGGAATCATAATTACTTTAACCTCCGGCTTA |
| core12 | ACGACAGTTGAGATTTAGGATCCTAATTTACGAGCAGCA |
| core13 | TAAATCATGATAAATAAGGCGTTATAT |
| core14 | TTACGAACAACATTATTACAATAATCGGCTGTCTTTTCG |
| core15 | TATAGTCCCTAAATTTAATGGTTAAGAC |
| core16 | GTTCAGCACACTATCATAACCCTGTAAAATGTTTA |
| core17 | AACAATACCAAAAGGAATTACGACTGCGGAATCGT |
| core18 | TATCCCAATACCACATTCAACTACCCTCAAATGCT |
| core19 | AACCAATCAGGTAGAAAGATTCATGAATGACCA |

| | |
|---|---|
| core20 | CATTCCAAGAATAAAACGAACTAACGCCTGACTAT |
| core21 | TTAGTTGCTATTTTCAAGAGTAATCTTGATTTGAA |
| core22 | TTGCGGGAGGTTATTACCCAAATCAACGGTCAA |
| core23 | AGAACGCGAGGCGTTCAGTGAATAAGGCTTCCATG |
| core24 | AATCAGATATAGAACCAGAACGAGTAGTACCTGAT |
| core25 | TAGGAATCATTACGGTTTAATTTCAACTTAAACAA |
| core26 | CCGGATATTCTTGAAGCCTTAAATCAAAACAACAT |
| core27 | AAGCTGCTCATTTTAGCGAACCTCCCCTGTTTATC |
| core28 | GACGAGAAACAGGCTTATCCGGTATTGAAAAATAA |
| core29 | GCTTGAGATCGCGCCCAATAGCAATGTAGA |
| core30 | ATTGTGAATTAGCCGTTTTTATTTTCACCTTAT |
| core31 | AGAGGACAGACCAAATAAGAAACGATACGCAGTAT |
| core32 | TCATAAGGGAAAATGAAAATAGCAAAAAGAAC |
| core33 | TTACTTAGCCTAACATAAAAACAGGGAACCGAGGA |
| core34 | AAATTGTGTCAGAATTAACTGAACACAAGCAGATA |
| core35 | AGTACAACGGAGGGTAATTGAGCGCTCTATCTTAC |
| core36 | TTGTTTAACGTCAAACCGAACTGACCAACCAAGAA |
| core37 | TTTACAGAGAGAAGGAACGAGGCGCAGACGTAACA |
| core38 | CGCATTAGACGGGGAAATCCGCGACCTGCTGCCCT |
| core39 | GAACAAAGTCAGAGATTTGTATCATCGAATTGG |
| core40 | ATCAGAGAGATAAGATTATACCAAGCGCGTAATC |
| core41 | GTTAGCACTTGCAGGGAGTTAAAGCGCCGACAATG |
| core42 | TGGCATGTCGTCACCCTCAGCAGAATTTCTTAA |
| core43 | AACGCAAGAACGAGGGTAGCAACATTGTATCGGTT |
| core44 | GCCGAACAGGACTAAAGACTTTTCTCCAAAAAAAA |
| core45 | CGAAGCCCATTAAACGGGTAAAATTGCGAATAATA |
| core46 | ACAACAGCGCCAAAGACAAACGCCTCCCTCAGA |
| core47 | AGTTGGCCGCTTTTGCGGGAATTAAGACTCCTTATTTTT |
| core48 | ACAGGATTGAGGGAGGGAAAAATCACCGGAA |
| core49 | AGGTGCGAAAGACAGCATCGTAATAACGGAATACCCGCC |
| core50 | TATCAATTATTCATTAAAGGCCTTATTAGCGTT |
| core51 | TTTAGGCTACAGAGGCTTTGAAAGTTACCAGAAGGAAAG |
| core52 | GGCTCCGACTTGAGCCATTTCGCGTTTTCATCG |
| core53 | AAATTCATGAGGAAGTTTCCTTTTTAAGAAAAGTCCT |
| core54 | ATTTCAAAATCACCAGTAGCAATCAAGTTTGCC |
| core55 | GGAATACGTAATGCCACTACAATGAAATAGCAATAGAAT |
| core56 | GCCCACCCTCATTTTCAGGGGTTTAGTACCGCCACGATT |
| core57 | CCGGAACAGGGCGACATTCAACCCTTGATACCGAT |
| core58 | CCAGGAACCCATGTACCGGCCCGGAATAGGTGTCTCA |
| core59 | ATAATCAGGTAAATATTGACGGAAGCTTGCTTTCG |
| core60 | TGCCACCAGTACAAACTACTAAGTGCCGTCGAGAGCTGA |
| core61 | ATAGCCCTGAATTATCACCGTCACCAAAAGGAGCC |
| core62 | GCACACAGACAGCCCTCATGATTAGCGGGGTTTTGGGCT |
| core63 | ACTGTAGGGGAATTAGAGCCAGTTTCACGTTGA |
| core64 | TTTCTAAAGTTTTGTCGTCAGAGGCTGAGACTCCTTAAG |
| core65 | GCGACAGACCATTACCATTAGCAGGAACAACTAAA |
| core66 | GTACTCAGGAGATAGCAAGCCCAATAGAGCCACCA |

| | |
|---|---|
| core67 | TATAAGTATATAACACTGAGTTTCGTCATCTTTTC |
| core68 | ACCAGGCGGAAACGCCTGTAGCATTCTTTTCGGTC |
| core69 | AAGGATTAGAGTTAGCGTAACGATAGCGTCAG |
| core70 | GAAAGTATTATTTCCAGACGTTAGTATAATCAGTA |
| core71 | GGCCTTGATATTCACAAACAAATAAATCATCACC |
| core72 | TTAAAGCCAGAATGGAAAGCGCAGTCTGGTTGA |
| core73 | ATTTACCGTTCCAGTAAGCGTCATACATCTCAGT |
| core74 | TTTGATGATACAGGAGTGTACTGGTAACAAGAG |
| core75 | TTTTAACGGGGTCAGTGCCTTGAGTAACAAACAT |

Tables S2

| Name | Sequence (5' - 3') |
|---|---|
| F-Core1 | TTGGGCGCCAGGGTGAACCCTAACCCTAACCCTAACCCAATGGTGGTTCCGAAAT |
| F-Core2 | TCCTGTGTGAAATTGAACCCTAACCCTAACCCTAACCCAAGCACGTATAACGTGC |
| F-Core3 | AAACGCTCATGGAAAAACCCTAACCCTAACCCTAACCCAATCAGGAAGATCGCAC |
| F-Core4 | GGCTGCGCAACTGTTAACCCTAACCCTAACCCTAACCCAAGCCTGAGTAGAAGAA |
| F-Core5 | CGTCGGATTCTCCGTAACCCTAACCCTAACCCTAACCCAATTTAATGCGCGAACT |
| F-Core6 | GAGCCGTCAATAGATAACCCTAACCCTAACCCTAACCCAACTACAAAGGCTATCA |
| F-Core7 | GTCAATCATATGTACAACCCTAACCCTAACCCTAACCCAAAGTTGGCAAATCAAC |
| F-Core8 | TGTGTAGGTAAAGATAACCCTAACCCTAACCCTAACCCAATGATGGCAATTCATC |
| F-Core9 | AAACATCAAGAAAACAACCCTAACCCTAACCCTAACCCAAGTTTGACCATTAGAT |
| F-Core10 | TGGCATCAATTCTACAACCCTAACCCTAACCCTAACCCAAGCTTTGAATACCAAG |
| F-Core11 | GTTTTTCTTTTCACCAAACCCTAACCCTAACCCTAACCCAAAAACGACGGCCAGTGC |
| F-Core12 | CCACACAACATACGAGAACCCTAACCCTAACCCTAACCCAACTACAGGGCGCGTACT |
| F-Core13 | TTATCCGCTCACAATTAACCCTAACCCTAACCCTAACCCAATGCGTTGCGCTCACTG |
| F-Core14 | CATGGTCATAGCTGTTAACCCTAACCCTAACCCTAACCCAAGGAAACCTGTCGTGCC |
| F-Core15 | AGCTCGAATTCGTAATAACCCTAACCCTAACCCTAACCCAAAGAGCGGGAGCTAAAC |
| F-Core16 | GGATCCCCGGGTACCGAACCCTAACCCTAACCCTAACCCAACGGCCAACGCGCGGGG |
| F-Core17 | CTGAGAAGTGTTTTTAAACCCTAACCCTAACCCTAACCCAAAGTAATAACATCACTT |
| F-Core18 | TAATCAGTGAGGCCACAACCCTAACCCTAACCCTAACCCAACAGTCACGACGTTGTA |
| F-Core19 | CTCAATCGTCTGAAATAACCCTAACCCTAACCCTAACCCAAGCCAGTTTGAGGGGAC |
| F-Core20 | TACCTACATTTTGACGAACCCTAACCCTAACCCTAACCCAAATGGTTGCTTTGACGA |
| F-Core21 | GCCATTGCAACAGGAAAACCCTAACCCTAACCCTAACCCAATTTCCTCGTTAGAATC |
| F-Core22 | AACAATATTACCGCCAAACCCTAACCCTAACCCTAACCCAAGCACCGCTTCTGGTGC |
| F-Core23 | TGCTGGTAATATCCAGAACCCTAACCCTAACCCTAACCCAAAGGAGGCCGATTAAAG |
| F-Core24 | GGGAAGGGCGATCGGTAACCCTAACCCTAACCCTAACCCAAATTTTTTAACCAATAG |
| F-Core25 | GCGGGCCTCTTCGCTAAACCCTAACCCTAACCCTAACCCAACAATACTTCTTTGATT |
| F-Core26 | TTGACCGTAATGGGATAACCCTAACCCTAACCCTAACCCAAGCACAGACAATATTTT |
| F-Core27 | GGGAACAAACGGCGGAAACCCTAACCCTAACCCTAACCCAAGACGACAGTATCGGCC |
| F-Core28 | TGAGCGAGTAACAACCAACCCTAACCCTAACCCTAACCCAATCCAGCCAGCTTTCCG |
| F-Core29 | TCATCAACATTAAATGAACCCTAACCCTAACCCTAACCCAACGCCATTAAAAATACC |
| F-Core30 | TTCCTGTAGCCAGCTTAACCCTAACCCTAACCCTAACCCAACGGAAACCAGGCAAAG |
| F-Core31 | TAACACCGCCTGCAACAACCCTAACCCTAACCCTAACCCAACAATCAATATCTGGTC |
| F-Core32 | AGTGCCACGCTGAGAGAACCCTAACCCTAACCCTAACCCAATTTGTTAAATCAGCTC |
| F-Core33 | TAGAAGTATTAGACTTAACCCTAACCCTAACCCTAACCCAAAATGCCGGAGAGGGTA |
| F-Core34 | AATACATTTGAGGATTAACCCTAACCCTAACCCTAACCCAATGAATGGCTATTAGTC |
| F-Core35 | TATCTTTAGGAGCACTAACCCTAACCCTAACCCTAACCCAATCTGGAGCAAACAAGA |
| F-Core36 | GGAAGGTTATCTAAAAAACCCTAACCCTAACCCTAACCCAAGAACGAACCACCAGCA |

| | |
|---|---|
| F-Core37 | CCCGGTTGATAATCAGAACCCTAACCCTAACCCTAACCCAAGACCCTGTAATACTTT |
| F-Core38 | AAAAGCCCCAAAAACAAACCCTAACCCTAACCCTAACCCAATCAAATATCAAACCCT |
| F-Core39 | GGCCGGAGACAGTCAAAACCCTAACCCTAACCCTAACCCAAGCGGAATTATCATCAT |
| F-Core40 | TCAAAAGGGTGAGAAAAACCCTAACCCTAACCCTAACCCAAGCTATTTTTGAGAGAT |
| F-Core41 | CCTCATATATTTAAAAACCCTAACCCTAACCCTAACCCAATTTGGATTATACTTCT |
| F-Core42 | TAAAAATTTTTAGAACAACCCTAACCCTAACCCTAACCCAAGAATCGATGAACGGTA |
| F-Core43 | AAAACAGAAATAAAGAAACCCTAACCCTAACCCTAACCCAAACGGATTCGCCTGATT |
| F-Core44 | AATTGCGTAGATTTTCAACCCTAACCCTAACCCTAACCCAAGTACCAAAAACATTAT |
| F-Core45 | AACAATTTCATTTGAAAACCCTAACCCTAACCCTAACCCAATTGATTCCCAATTCTG |
| F-Core46 | AAAATTAATTACATTTAACCCTAACCCTAACCCTAACCCAAATTCCTGATTATCAGA |
| F-Core47 | AAGAAGATGATGAAACAACCCTAACCCTAACCCTAACCCAAAATATAATCCTGATTG |
| F-Core48 | TCAATTACCTGAGCAAAACCCTAACCCTAACCCTAACCCAACAATAACCTGTTTAGC |
| F-Core49 | AGGCGAATTATTCATTAACCCTAACCCTAACCCTAACCCAAGAATAATGGAAGGGTT |
| F-Core50 | TAATAGTAGTAGCATTAACCCTAACCCTAACCCTAACCCAACGAACCAGACCGGAAG |
| F-Core51 | AACATCCAATAAATCAAACCCTAACCCTAACCCTAACCCAAATCGGGAGAAACAATA |
| F-Core52 | ATAATGCTGTAGCTCAAACCCTAACCCTAACCCTAACCCAACGAACGAGTAGATTTA |
| F-Core53 | TTGCGGATGGCTTAGAAACCCTAACCCTAACCCTAACCCAAACATTTCGCAAATGGT |
| F-Core54 | CTTTAATTGCTCCTTTAACCCTAACCCTAACCCTAACCCAATATATTTTCATTTGGG |
| F-Core55 | GGGCGAACCCTAACCCTAACCCTAACCCAAGCTAACTCACATTAAT |
| F-Core56 | CACTAAACCCTAACCCTAACCCTAACCCAACCCGCTTTCCAGTCG |
| F-Core57 | TTGAGAACCCTAACCCTAACCCTAACCCAAAGCTGCATTAATGAAT |
| F-Core58 | CGGCAAAATCCCTTATAACCCTAACCCTAACCCTAACCCAAAGAGGCGGTT |
| F-Core59 | GTGAGACGGGCAACAGAACCCTAACCCTAACCCTAACCCAAGGTTTG |
| F-Core60 | TAGACAGGAAAACCCTAACCCTAACCCTAACCCAACTCAAACT |
| F-Core61 | ATCGGCCTAACCCTAACCCTAACCCTAACCCAACGCCATTCGC |
| F-Core62 | AAAACAGAGGAACCCTAACCCTAACCCTAACCCAAAGTTGAAA |
| F-Core63 | GGAATTGAAACCCTAACCCTAACCCTAACCCAAATCGTAAAAC |
| F-Core64 | TACCATATCAAACCCTAACCCTAACCCTAACCCAATTACAAAA |
| F-Core65 | TCGCGCAGAACCCTAACCCTAACCCTAACCCAAGCGCGAGCTG |
| F-Core66 | ACATGTTTTAAATATGAACCCTAACCCTAACCCTAACCCAACATAGCGATAGC |
| F-Core67 | GCTTAATTGCTGAATAACCCTAACCCTAACCCTAACCCAAGAGTCAATAGT |
| F-Core68 | TGATAAGAGGTCATTTAACCCTAACCCTAACCCTAACCCAATCTGAGAGACTA |
| F-Core69 | AAAGAACGCGAAACCCTAACCCTAACCCTAACCCAAAATTCGAGCTTCAAAG |
| F-Core70 | TTAAAGAACGTGGACTCCAACGTCAAA |
| F-Core71 | TGTTGTTCCAGTTTGGAACAAGAGTC |
| F-Core72 | AAATCAAAAGAATAGCCCGAGATAGGG |
| F-Core73 | CCCCAGCAGGCGAAAATCCTGTTTGA |
| Cy3-I38 | Cy3-AACAACTAATAGATTAAACCCTAACCCTAACCCTAACCCAAGATAGCCCTAAAACAT |
| I45-Cy5 | TGCAATGCCTGAGTAAAACCCTAACCCTAACCCTAACCCAAGGTCATTGCCTGAGAG-Cy5 |
| I1-Biotin | AAAAGCAGGTCGACTCTAGAAACCCTAACCCTAACCCTAACCCAAGGATTT-Biotin |
| I2-Biotin | AAAATAATTCGCGTCTGGCCAACCCTAACCCTAACCCTAACCCAAGAAGAT-Biotin |
| I3-Biotin | AAAATATTTCAACGCAAGGAAACCCTAACCCTAACCCTAACCCAAAGAACC-Biotin |
| I4-Biotin | AAAACAGGATTAGAGAGTACAACCCTAACCCTAACCCTAACCCAAGGTTGGGTTATA-Biotin |

Tables S3

| Name | Sequence (5' - 3') |
|---|---|
| F-Core1 | TTGGGCGCCAGGGTGTGCGACCGTTTTTTTTACGGTCGCTTGGTGGTTCCGAAAT |
| F-Core2 | TCCTGTGTGAAATTGTGCGACCGTTTTTTTTACGGTCGCTGCACGTATAACGTGC |

| | |
|---|---|
| F-Core3 | AAACGCTCATGGAAATGCGACCGTTTTTTTTACGGTCGCTTCAGGAAGATCGCAC |
| F-Core4 | GGCTGCGCAACTGTTTGCGACCGTTTTTTTTACGGTCGCTGCCTGAGTAGAAGAA |
| F-Core5 | CGTCGGATTCTCCGTTGCGACCGTTTTTTTTACGGTCGCTTTTAATGCGCGAACT |
| F-Core6 | GAGCCGTCAATAGATTGCGACCGTTTTTTTTACGGTCGCTCTACAAAGGCTATCA |
| F-Core7 | GTCAATCATATGTACTGCGACCGTTTTTTTTACGGTCGCTAGTTGGCAAATCAAC |
| F-Core8 | TGTGTAGGTAAAGATTGCGACCGTTTTTTTTACGGTCGCTTGATGGCAATTCATC |
| F-Core9 | AAACATCAAGAAAACTGCGACCGTTTTTTTTACGGTCGCTGTTTGACCATTAGAT |
| F-Core10 | TGGCATCAATTCTACTGCGACCGTTTTTTTTACGGTCGCTGCTTTGAATACCAAG |
| F-Core11 | GTTTTTCTTTTCACCATGCGACCGTTTTTTTTACGGTCGCTAAACGACGGCCAGTGC |
| F-Core12 | CCACACAACATACGAGTGCGACCGTTTTTTTTACGGTCGCTCTACAGGGCGCGTACT |
| F-Core13 | TTATCCGCTCACAATTTGCGACCGTTTTTTTTACGGTCGCTTGCGTTGCGCTCACTG |
| F-Core14 | CATGGTCATAGCTGTTTGCGACCGTTTTTTTTACGGTCGCTGGAAACCTGTCGTGCC |
| F-Core15 | AGCTCGAATTCGTAATTGCGACCGTTTTTTTTACGGTCGCTAGAGCGGGAGCTAAAC |
| F-Core16 | GGATCCCCGGGTACCGTGCGACCGTTTTTTTTACGGTCGCTCGGCCAACGCGCGGGG |
| F-Core17 | CTGAGAAGTGTTTTTATGCGACCGTTTTTTTTACGGTCGCTAGTAATAACATCACTT |
| F-Core18 | TAATCAGTGAGGCCACTGCGACCGTTTTTTTTACGGTCGCTCAGTCACGACGTTGTA |
| F-Core19 | CTCAATCGTCTGAAATTGCGACCGTTTTTTTTACGGTCGCTGCCAGTTTGAGGGGAC |
| F-Core20 | TACCTACATTTTGACGTGCGACCGTTTTTTTTACGGTCGCTATGGTTGCTTTGACGA |
| F-Core21 | GCCATTGCAACAGGAATGCGACCGTTTTTTTTACGGTCGCTTTTCCTCGTTAGAATC |
| F-Core22 | AACAATATTACCGCCATGCGACCGTTTTTTTTACGGTCGCTGCACCGCTTCTGGTGC |
| F-Core23 | TGCTGGTAATATCCAGTGCGACCGTTTTTTTTACGGTCGCTAGGAGGCCGATTAAAG |
| F-Core24 | GGGAAGGGCGATCGGTTGCGACCGTTTTTTTTACGGTCGCTATTTTTTAACCAATAG |
| F-Core25 | GCGGGCCTCTTCGCTATGCGACCGTTTTTTTTACGGTCGCTCAATACTTCTTTGATT |
| F-Core26 | TTGACCGTAATGGGATTGCGACCGTTTTTTTTACGGTCGCTGCACAGACAATATTTT |
| F-Core27 | GGGAACAAACGGCGGATGCGACCGTTTTTTTTACGGTCGCTGACGACAGTATCGGCC |
| F-Core28 | TGAGCGAGTAACAACCTGCGACCGTTTTTTTTACGGTCGCTTCCAGCCAGCTTTCCG |
| F-Core29 | TCATCAACATTAAATGTGCGACCGTTTTTTTTACGGTCGCTCGCCATTAAAAATACC |
| F-Core30 | TTCCTGTAGCCAGCTTTGCGACCGTTTTTTTTACGGTCGCTCGGAAACCAGGCAAAG |
| F-Core31 | TAACACCGCCTGCAACTGCGACCGTTTTTTTTACGGTCGCTCAATCAATATCTGGTC |
| F-Core32 | AGTGCCACGCTGAGAGTGCGACCGTTTTTTTTACGGTCGCTTTTGTTAAATCAGCTC |
| F-Core33 | TAGAAGTATTAGACTTTGCGACCGTTTTTTTTACGGTCGCTAATGCCGGAGAGGGTA |
| F-Core34 | AATACATTTGAGGATTTGCGACCGTTTTTTTTACGGTCGCTTGAATGGCTATTAGTC |
| F-Core35 | TATCTTTAGGAGCACTTGCGACCGTTTTTTTTACGGTCGCTTCTGGAGCAAACAAGA |
| F-Core36 | GGAAGGTTATCTAAAATGCGACCGTTTTTTTTACGGTCGCTGAACGAACCACCAGCA |
| F-Core37 | CCCGGTTGATAATCAGTGCGACCGTTTTTTTTACGGTCGCTGACCCTGTAATACTTT |
| F-Core38 | AAAAGCCCCAAAAACATGCGACCGTTTTTTTTACGGTCGCTTCAAATATCAAACCCT |
| F-Core39 | GGCCGGAGACAGTCAATGCGACCGTTTTTTTTACGGTCGCTGCGGAATTATCATCAT |
| F-Core40 | TCAAAAGGGTGAGAAATGCGACCGTTTTTTTTACGGTCGCTGCTATTTTTGAGAGAT |
| F-Core41 | CCTCATATATTTTAAATGCGACCGTTTTTTTTACGGTCGCTTTTGGATTATACTTCT |
| F-Core42 | TAAAAATTTTTAGAACTGCGACCGTTTTTTTTACGGTCGCTGAATCGATGAACGGTA |
| F-Core43 | AAAACAGAAATAAAGATGCGACCGTTTTTTTTACGGTCGCTACGGATTCGCCTGATT |
| F-Core44 | AATTGCGTAGATTTTCTGCGACCGTTTTTTTTACGGTCGCTGTACCAAAAACATTAT |
| F-Core45 | AACAATTTCATTTGAATGCGACCGTTTTTTTTACGGTCGCTTTGATTCCCAATTCTG |
| F-Core46 | AAAATTAATTACATTTTGCGACCGTTTTTTTTACGGTCGCTATTCCTGATTATCAGA |
| F-Core47 | AAGAAGATGATGAAACTGCGACCGTTTTTTTTACGGTCGCTAATATAATCCTGATTG |
| F-Core48 | TCAATTACCTGAGCAATGCGACCGTTTTTTTTACGGTCGCTCAATAACCTGTTTAGC |
| F-Core49 | AGGCGAATTATTCATTTGCGACCGTTTTTTTTACGGTCGCTGAATAATGGAAGGGTT |

| | |
|---|---|
| F-Core50 | TAATAGTAGTAGCATTTGCGACCGTTTTTTTTACGGTCGCTCGAACCAGACCGGAAG |
| F-Core51 | AACATCCAATAAATCATGCGACCGTTTTTTTTACGGTCGCTATCGGGAGAAACAATA |
| F-Core52 | ATAATGCTGTAGCTCATGCGACCGTTTTTTTTACGGTCGCTCGAACGAGTAGATTTA |
| F-Core53 | TTGCGGATGGCTTAGATGCGACCGTTTTTTTTACGGTCGCTACATTTCGCAAATGGT |
| F-Core54 | CTTTAATTGCTCCTTTTGCGACCGTTTTTTTTACGGTCGCTTATATTTTCATTTGGG |
| F-Core55 | GGGCGTGCGACCGTTTTTTTTACGGTCGCTGCTAACTCACATTAAT |
| F-Core56 | CACTATGCGACCGTTTTTTTTACGGTCGCTCCCGCTTTCCAGTCG |
| F-Core57 | TTGAGTGCGACCGTTTTTTTTACGGTCGCTAGCTGCATTAATGAAT |
| F-Core58 | CGGCAAAATCCCTTATTGCGACCGTTTTTTTTACGGTCGCTAGAGGCGGTT |
| F-Core59 | GTGAGACGGGCAACAGTGCGACCGTTTTTTTTACGGTCGCTGGTTTG |
| F-Core60 | TAGACAGGAATGCGACCGTTTTTTTTACGGTCGCTCTCAAACT |
| F-Core61 | ATCGGCCTTGCGACCGTTTTTTTTACGGTCGCTCGCCATTCGC |
| F-Core62 | AAAACAGAGGTGCGACCGTTTTTTTTACGGTCGCTAGTTGAAA |
| F-Core63 | GGAATTGATGCGACCGTTTTTTTTACGGTCGCTATCGTAAAAC |
| F-Core64 | TACCATATCATGCGACCGTTTTTTTTACGGTCGCTTTACAAAA |
| F-Core65 | TCGCGCAGTGCGACCGTTTTTTTTACGGTCGCTGCGCGAGCTG |
| F-Core66 | ACATGTTTTAAATATGTGCGACCGTTTTTTTTACGGTCGCTCATAGCGATAGC |
| F-Core67 | GCTTAATTGCTGAATTGCGACCGTTTTTTTTACGGTCGCTGAGTCAATAGT |
| F-Core68 | TGATAAGAGGTCATTTTGCGACCGTTTTTTTTACGGTCGCTTCTGAGAGACTA |
| F-Core69 | AAAGAACGCGATGCGACCGTTTTTTTTACGGTCGCTAATTCGAGCTTCAAAG |
| F-Core70 | TTAAAGAACGTGGACTCCAACGTCAAA |
| F-Core71 | TGTTGTTCCAGTTTGGAACAAGAGTC |
| F-Core72 | AAATCAAAAGAATAGCCCGAGATAGGG |
| F-Core73 | CCCCAGCAGGCGAAAATCCTGTTTGA |
| Cy3-H38 | Cy3-AACAACTAATAGATTATGCGACCGTTTTTTTTACGGTCGCTGATAGCCCTAAAACAT |
| H45-Cy5 | TGCAATGCCTGAGTAATGCGACCGTTTTTTTTACGGTCGCTGGTCATTGCCTGAGAG-Cy5 |
| H1-Biotin | AAAAGCAGGTCGACTCTAGATGCGACCGTTTTTTTTACGGTCGCTGGATTT-Biotin |
| H2-Biotin | AAAATAATTCGCGTCTGGCCTGCGACCGTTTTTTTTACGGTCGCTGAAGAT-Biotin |
| H3-Biotin | AAAATATTTCAACGCAAGGATGCGACCGTTTTTTTTACGGTCGCTAGAACC-Biotin |
| H4-Biotin | AAAACAGGATTAGAGAGTACTGCGACCGTTTTTTTTACGGTCGCTGGTTGGGTTATA-Biotin |

Tables S4

| Name | Sequence (5' - 3') |
|---|---|
| hairpin-a | AGCGACCGTAAAAAAAACGGTCGCAAAAAAAAAA |
| hairpin-b | TTTTTTTTTTGCGACCGTTTTTTTTACGGTCGCT |
| i-motif-a | TTTTTTTTGTGTTAGTGTTAGTGTTAGTGTT |
| i-motif-b | AACACTAACACTAACACTAACACAAAAAAAA |

#### 3 Supporting figures

##### 3.1 figures S1-S5

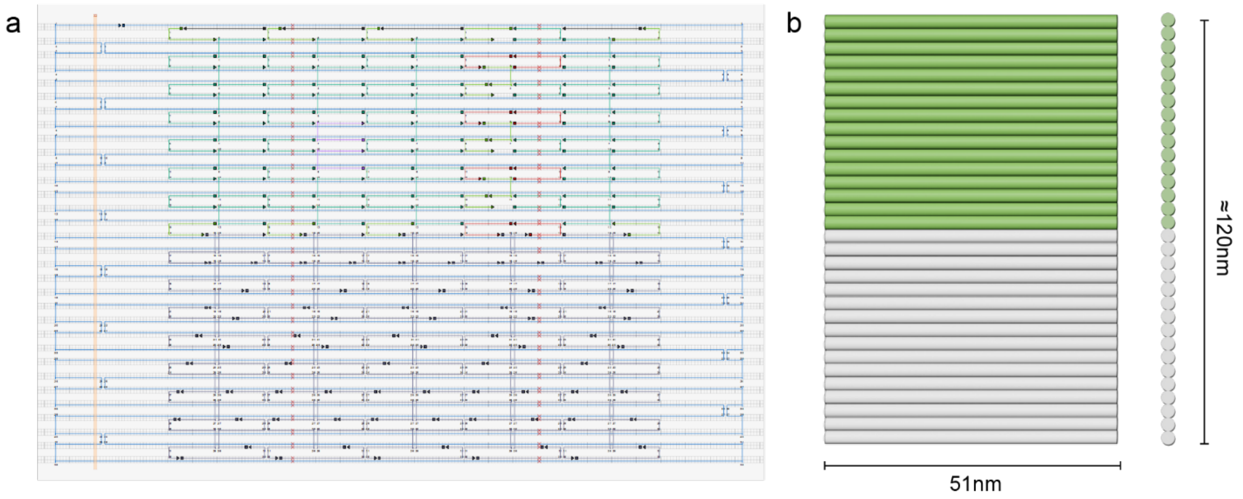


Figure S1. Design of DNA origami nanostructures. (a) DNA origami structure designed according to caDNAno software. The structure consists of 32 DNA double helices, each containing 150 bases. The upper half is a crossover replaced scalable structure and the lower half is a normal DNA origami structure. Sequence information is shown in Tables S1-S4.(b) Front and side views of the DNA origami nanostructures, the lateral width of the double helix part of the structure is 51 nm and the longitudinal length is about 120 nm.

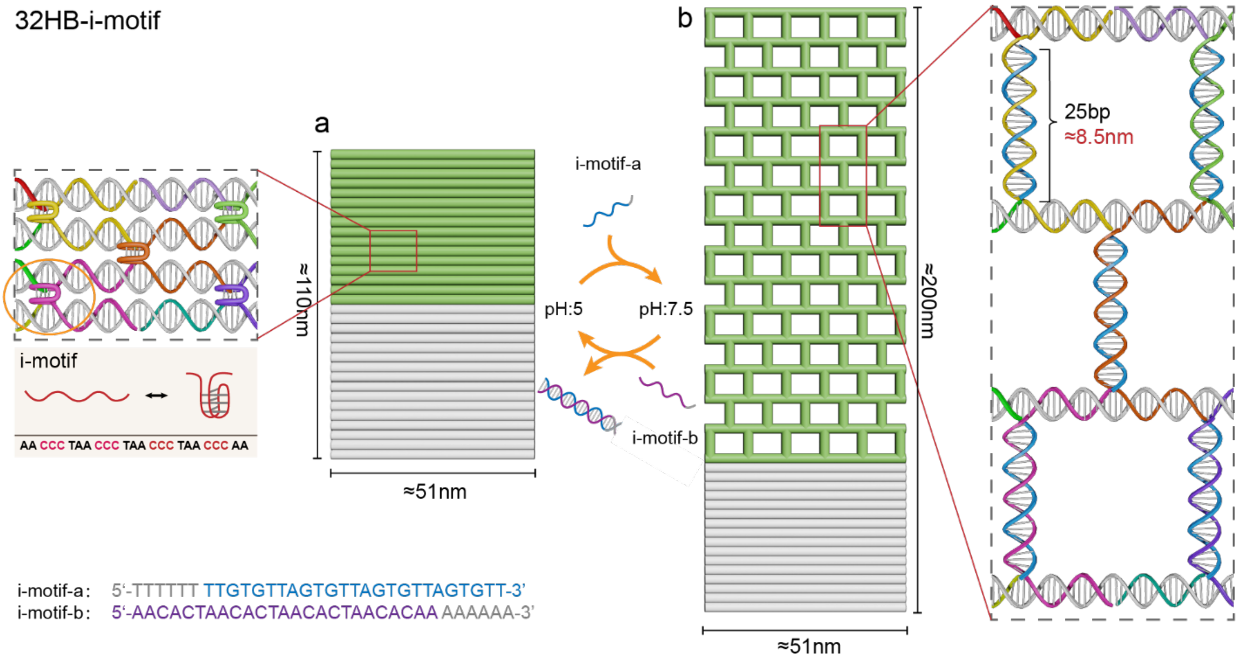


Figure S2. Schematic representation of the 32HB-i-motif in contraction. (a) Schematic representation of 32HB-i-motif in a contracted state at low ph. During the elevation of pH=5 to pH=7.5, the i-motif stretches and is complemented by the complementary chain i-motif-a, and 32HB-i-motif transforms to a stretched state. During the reduction of pH=7.5 to pH=5, the i-motif contracts and is complemented by the complementary strand i-motif-b to the i-motif-a as a DNA double helix, and 32HB-i-motif transforms to the contracted state. (b) Schematic representation of 32HB-i-motif in a stretched state at high pH.

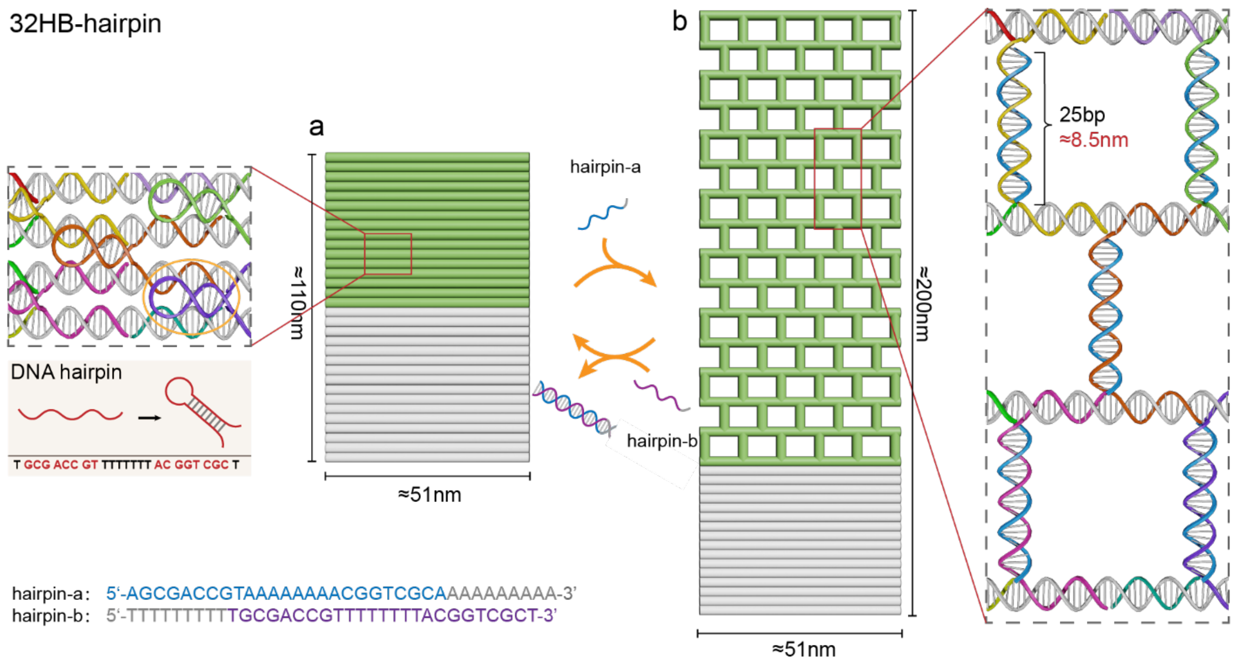


Figure S3. Schematic representation of the 32HB-hairpin in a contracted state. (a) Schematic representation of the contracted state of 32HB-hairpin. Upon addition of the complementary strand hairlpin-a, the DNA-hairpin is complemented to a double helix and the 32HB-hairpin transforms to a stretched state. Addition of complementary strand hairlpin-b replaces hairlpin-a by complementation, and 32HB-hairpin transforms to a contracted state. (b) Schematic representation of 32HB-hairpin in the stretched state.

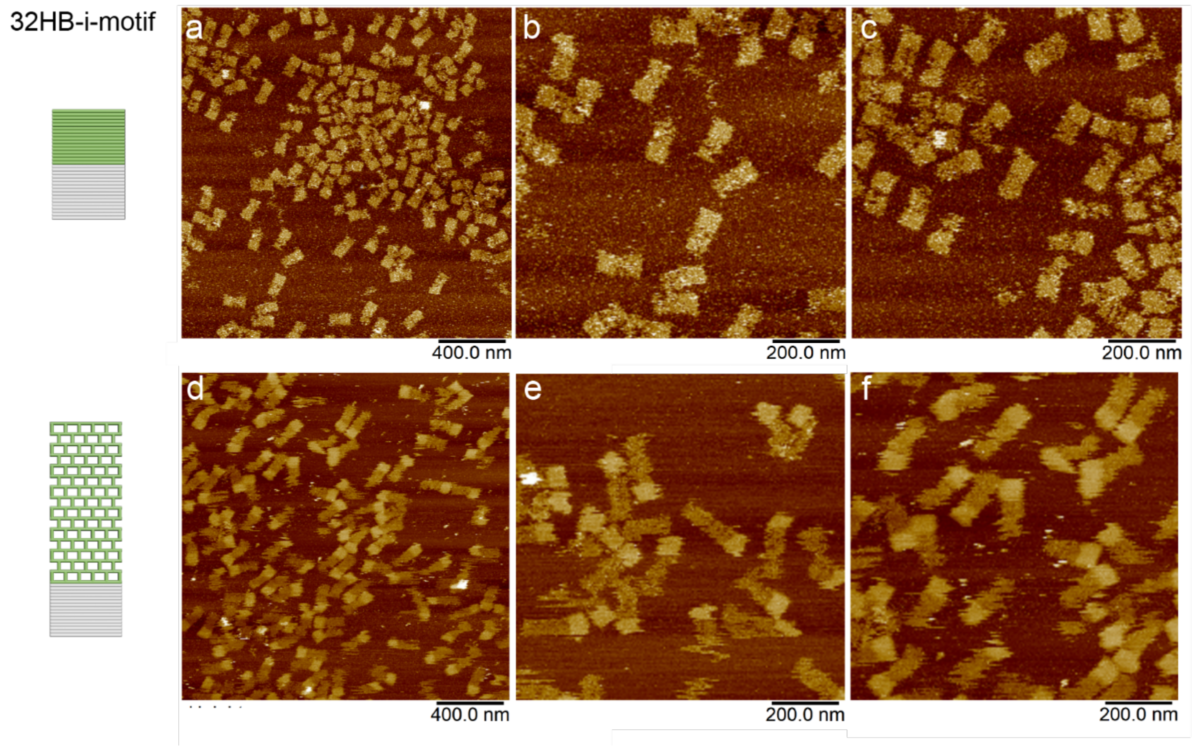


Figure S4. AFM images of 32HB-i-motif. (a-c) AFM images of 32HB-i-motif in contracted state. (b-f) AFM images of 32HB-i-motif in the stretched state. Scale bars: (a, d): 400 nm; (b,c,e,f): 200 nm.

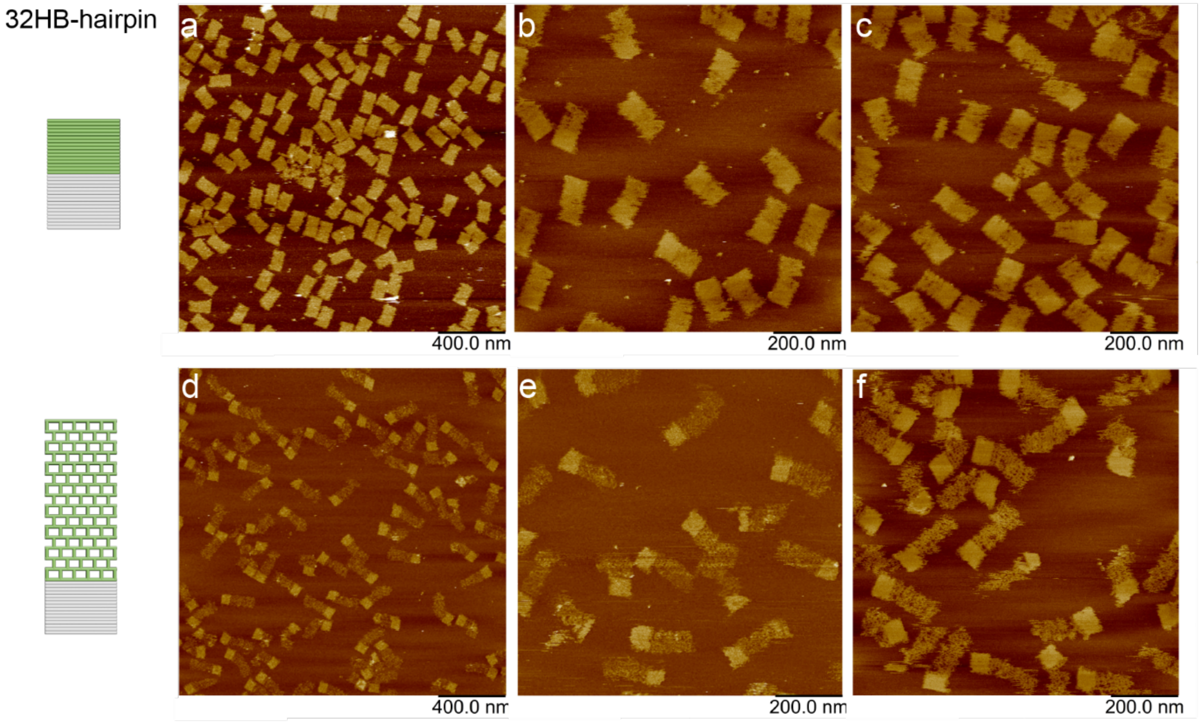


Figure S5. AFM images of 32HB-hairpin. (a-c) AFM images of 32HB-hairpin in contracted state. (b-f) AFM images of 32HB-hairpin in the stretched state. Scale bars: (a, d): 400 nm; (b,c,e,f): 200 nm.
